# Hybrid decay: a transgenerational epigenetic decline in vigor and viability triggered in backcross populations of teosinte with maize

**DOI:** 10.1101/588715

**Authors:** Wei Xue, Sarah N. Anderson, Xufeng Wang, Liyan Yang, Peter A. Crisp, Qing Li, Jaclyn Noshay, Patrice S. Albert, James A. Birchler, Paul Bilinski, Michelle C. Stitzer, Jeffrey Ross-Ibarra, Sherry Flint-Garcia, Xuemei Chen, Nathan M. Springer, John F. Doebley

**Affiliations:** Department of Genetics, University of Wisconsin, Madison, Wisconsin 53706; College of Agronomy, Shenyang Agricultural University, 110866, Liaoning Province, China; Department of Plant and Microbial Biology, University of Minnesota, St. Paul, Minnesota 55108; Guangdong Provincial Key Laboratory for Plant Epigenetics, Shenzhen University, 518060, Guangdong Province, China; Life Science College, Shanxi Normal University, 041004, Shanxi Province, China; Division of Biological Sciences, University of Missouri, Columbia, Missouri 65211; Department of Plant Sciences, University of California, Davis, California 95616; Agricultural Research Service, United States Department of Agriculture, Columbia, Missouri, 65211; Department of Botany and Plant Sciences, University of California, Riverside, California 92521

**Keywords:** Zea mays, maize, teosinte, epigenetic, transposable element, CNVs, sRNAs

## Abstract

In the course of generating populations of maize with teosinte chromosomal introgressions, an unusual sickly plant phenotype was noted in individuals from crosses with two teosinte accessions collected near Valle de Bravo, Mexico. The plants of these Bravo teosinte accessions appear phenotypically normal themselves and the F_1_ plants appear similar to typical maize x teosinte F_1_s. However, upon backcrossing to maize, the BC_1_ and subsequent generations display a number of detrimental characteristics including shorter stature, reduced seed set and abnormal floral structures. This phenomenon is observed in all BC individuals and there is no chromosomal segment linked to the sickly plant phenotype in advanced backcross generations. Once the sickly phenotype appears in a lineage, normal plants are never again recovered by continued backcrossing to the normal maize parent. Whole-genome shotgun sequencing reveals a small number of genomic sequences, some with homology to transposable elements, that have increased in copy number in the backcross populations. Transcriptome analysis of seedlings, which do not have striking phenotypic abnormalities, identified segments of 18 maize genes that exhibit increased expression in sickly plants. A *de novo* assembly of transcripts present in plants exhibiting the sickly phenotype identified a set of 59 up-regulated novel transcripts. These transcripts include some examples with sequence similarity to transposable elements and other sequences present in the recurrent maize parent (W22) genome as well as novel sequences not present in the W22 genome. Genome-wide profiles of gene expression, DNA methylation and sRNAs are similar between sickly plants and normal controls, although a few up-regulated transcripts and transposable elements are associated with altered sRNA or methylation profiles. This study documents hybrid incompatibility and genome instability triggered by the backcrossing of Bravo teosinte with maize. We name this phenomenon “hybrid decay” and present ideas on the mechanism that may underlie it.

## ARTICLE SUMMARY

We describe a phenomenon in maize and its nearest wild relative, teosinte, by which backcross progeny of a specific teosinte and maize exhibit a sickly whole-plant phenotype involving changes in morphology, vigor and viability. We characterized the sickly backcross plants and normal control plants by multiple assays including mRNA expression, sRNA expression, genome-wide methylation patterns, karyotype and changes in genome content. Diverse sequences (some with homology to transposable elements) are increased in copy number and expression and there is elevated abundance of sRNAs with homology to these sequences. We call the phenomenon “hybrid decay” and discuss potential underlying mechanisms.

Plant breeders seek to develop improved varieties through crosses among different individuals of a species followed by selection. New combinations of alleles can lead to improved performance, allowing development of elite varieties. In many plant species, the direct combination of genetic information in the two parents can lead to hybrid vigor (heterosis). In other cases, there can be deleterious consequences of crossing individuals that is often referred to as hybrid incompatibility. This phenomenon has been particularly well-studied for crosses between members of related species (Bomblies and Weigel 2007; Rieseberg and Blackman 2010; Fishman and Sweigart 2018). Even within a species there are examples of combinations that can lead to reduced vigor or fertility (Bomblies 2010). In some classic examples, hybrid incompatibility is caused by chromosomal rearrangements such as inversions or translocations that can result in partial sterility (Fishman and Sweigart 2018).

Maize geneticists have a long history of crossing maize (*Zea mays* ssp. *mays*) and its wild relatives, the annual Mexican teosintes (*Zea mays* ssp. *parviglumis* or *Zea mays* ssp. *mexicana*) for a variety of reasons. Beadle studied chromosome pairing in maize-teosinte hybrids and the inheritance of domestication traits in a maize-teosinte F_2_ population (Beadle 1932, 1972). Kermicle studied pollen-pistil incompatibility in maize-teosinte hybrids and their derivatives (Kermicle 2006). Multiple QTL studies have mapped the genes controlling domestication traits in maize-teosinte hybrid populations (e. g. Doebley and Stec 1991; Briggs *et al.* 2007). Other studies have utilized populations derived from maize x teosinte crosses to map and identify key domestication genes including *teosinte branched 1*(*tb1*), *teosinte glume architecture* (*tga1*), and *prolificacy1.1* (*prol1.1*) (Doebley *et al.* 1997; Wang *et al.* 2005; Wills *et al.* 2013). Finally, two projects assayed the effects of teosinte chromosome segments introgressed into maize on a variety of domestication and agronomic traits (Studer and Doebley 2012; Liu *et al.* 2016). Although there are a few known chromosomal inversions that are polymorphic in teosinte and maize (Fang *et al.* 2012) and a few polymorphic factors for pollen-pistil compatibility that can prevent hybrid formation (Lu *et al.* 2014), there are no reports of a severe loss of vigor or viability among maize-teosinte hybrids or their descendant lines.

During the construction mapping populations to study the effects of teosinte alleles, we noted an unusual sickly syndrome in the backcross progeny resulting from crosses of maize with teosinte from near the Valle de Bravo in the state of Mexico, hereafter “Bravo” teosinte. The initial F_1_ hybrids are normal, but a sickly phenotype is observed in the Backcross 1 (BC_1_) and is more pronounced in subsequent backcross generations. Once the sickly phenotype appears in a lineage, healthy plants are never recovered by additional backcrosses to the normal maize parent. The inheritance pattern for the hybrid decay syndrome does not depend upon inheritance of specific chromosomal segments from the teosinte parent and does not segregate in backcross populations. We documented genomic instability in these backcross populations with increased copy number for specific sequences some with homology to transposable elements. *De novo* assembly of transcripts identifies a collection of up-regulated sequences including some that have little or no similarity to sequences in the genomes of the maize recurrent parents (W22 or B73). Although the global patterns of DNA methylation and sRNA production are similar between sickly and normal plants, some transposable elements and sequences that are up-regulated in sickly plants show altered methylation and sRNA profiles. Our observations suggest that crosses between Bravo teosinte and maize can trigger genomic instability that is inherited in all progeny. We name this phenomenon “hybrid decay” - a transgenerational decline in vigor and viability triggered in backcross populations of Bravo teosinte with maize.

## Results

### Discovery of a hybrid decay phenomenon in maize x Bravo populations

Two projects assayed the effects of teosinte chromosomal segments that had been transferred into maize by backcross breeding (Studer and Doebley 2012; Liu *et al.* 2016). These two projects used teosinte accessions collected from various geographic locations (Figure S1). Two of the accessions of *Zea mays* ssp. *parviglumis* came from near Valle de Bravo in the state of Mexico. These two accessions have typical teosinte characteristics (Figure S2); however, observations from these projects revealed that the crosses of maize lines with teosinte from the Valle de Bravo region result in a loss of vigor and fertility in backcross generations. We use the term “sickly syndrome” for the collection of abnormal traits that appear in the backcross plants and the term “hybrid decay” for this phenomenon and its unknown causal mechanism. While F_1_ hybrids of maize and Bravo teosinte are phenotypically normal, some sickly plants appear in the BC_1_, and all plants in the BC_2_ and later backcross generations are sickly. We did not observe a notable increase in severity beyond the BC_2_ generation.

The initial observation of hybrid decay was made during a project assessing the phenotypic effects of allelic diversity within teosinte for the domestication gene *teosinte branched 1*(*tb1*) (Studer and Doebley 2012). Ten accessions of teosinte were crossed and then backcrossed into W22 (recurrent and female parent) for multiple generations, including five *Zea mays* ssp*. parviglumis*, four *Zea mays* ssp*. mexicana*, and one *Zea diploperennis* (Figure 1A). The F_1_ plants of all crosses exhibited phenotypes typical of maize x teosinte F_1_ hybrids. The backcross populations from all teosinte accessions other than the Bravo teosinte accession exhibited the expected variation in morphological traits that distinguish maize and teosinte, but the plants of these populations were otherwise healthy and produced viable offspring. However, in crosses of the Bravo teosinte accession (Beadle and Kato Site 6 from 79 km south of Valle de Bravo), multiple phenotypic abnormalities including reduced stature, abnormal floral morphology and reduced fertility were observed in the BC_2_ and subsequent backcross generations (Figure 1A, Figure 2). Moreover, all backcross plants from multiple independent single seed descent backcross lineages derived from the Bravo accession exhibit this sickly syndrome, suggesting the phenotype is not due to a single chromosomal region from Bravo segregating in a Mendelian fashion.

**Figure 1.**
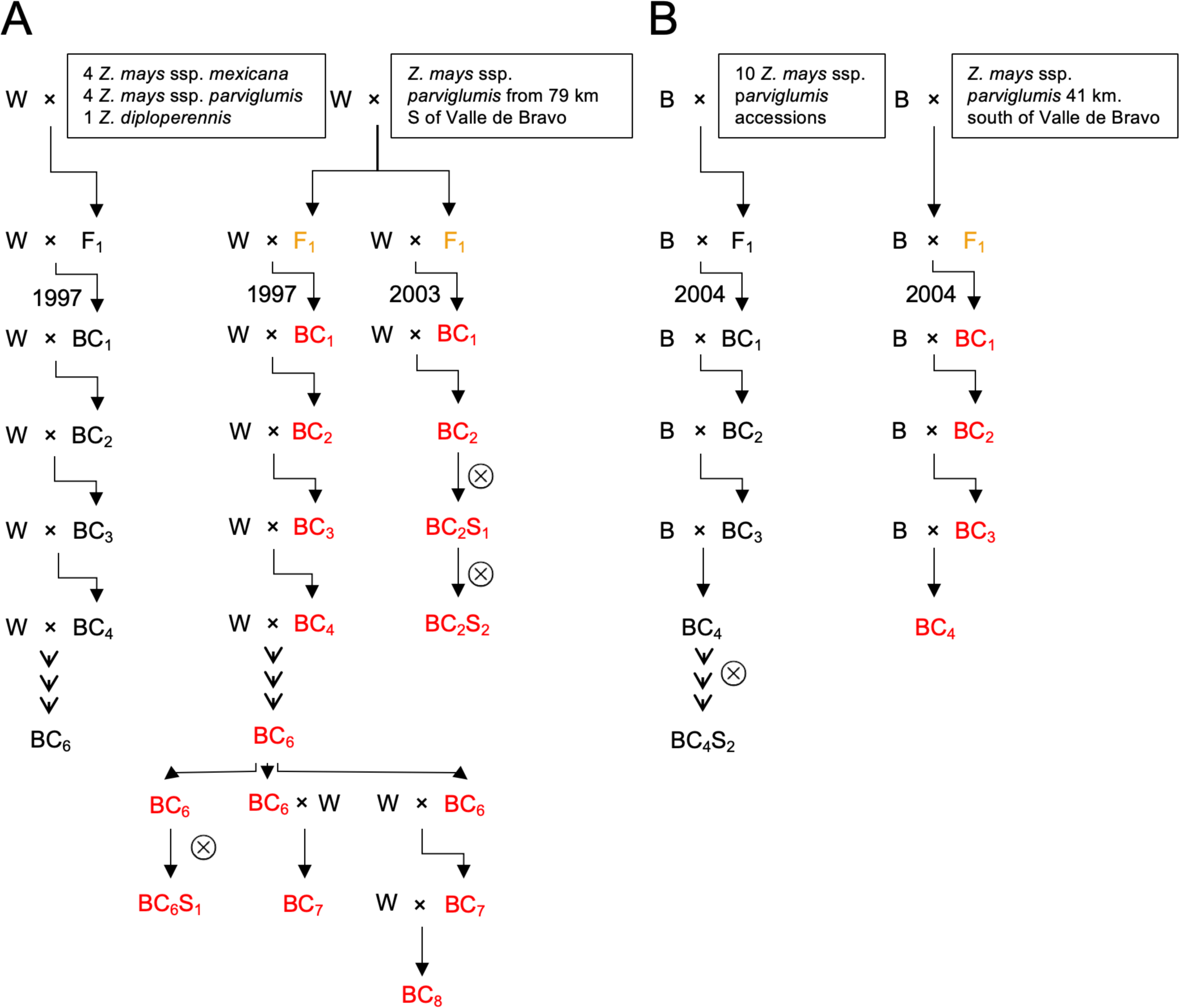
Origins of the hybrid decay syndrome within the pedigrees of materials produced in this study. The crossing scheme is shown to illustrate the pedigree of the materials and the appearance of the hybrid decay syndrome. The crossing schemes utilized in the Doebley lab (A) and Flint-Garcia lab (B) are shown. The female parent of each cross is listed first (W indicates W22 and B indicates B73 in the pedigrees). The left portions of panels A and B illustrate the crossing scheme used for multiple other accessions as a single pedigree and in all cases these individuals did not display the phenotypic abnormalities of hybrid decay. Any generations that exhibit the appearance of the hybrid decay syndrome are shown using red text. The generations that have progeny in subsequent generations that exhibit the hybrid decay syndrome are shown using orange text but were not carefully assessed for phenotypes.

**Figure 2.**
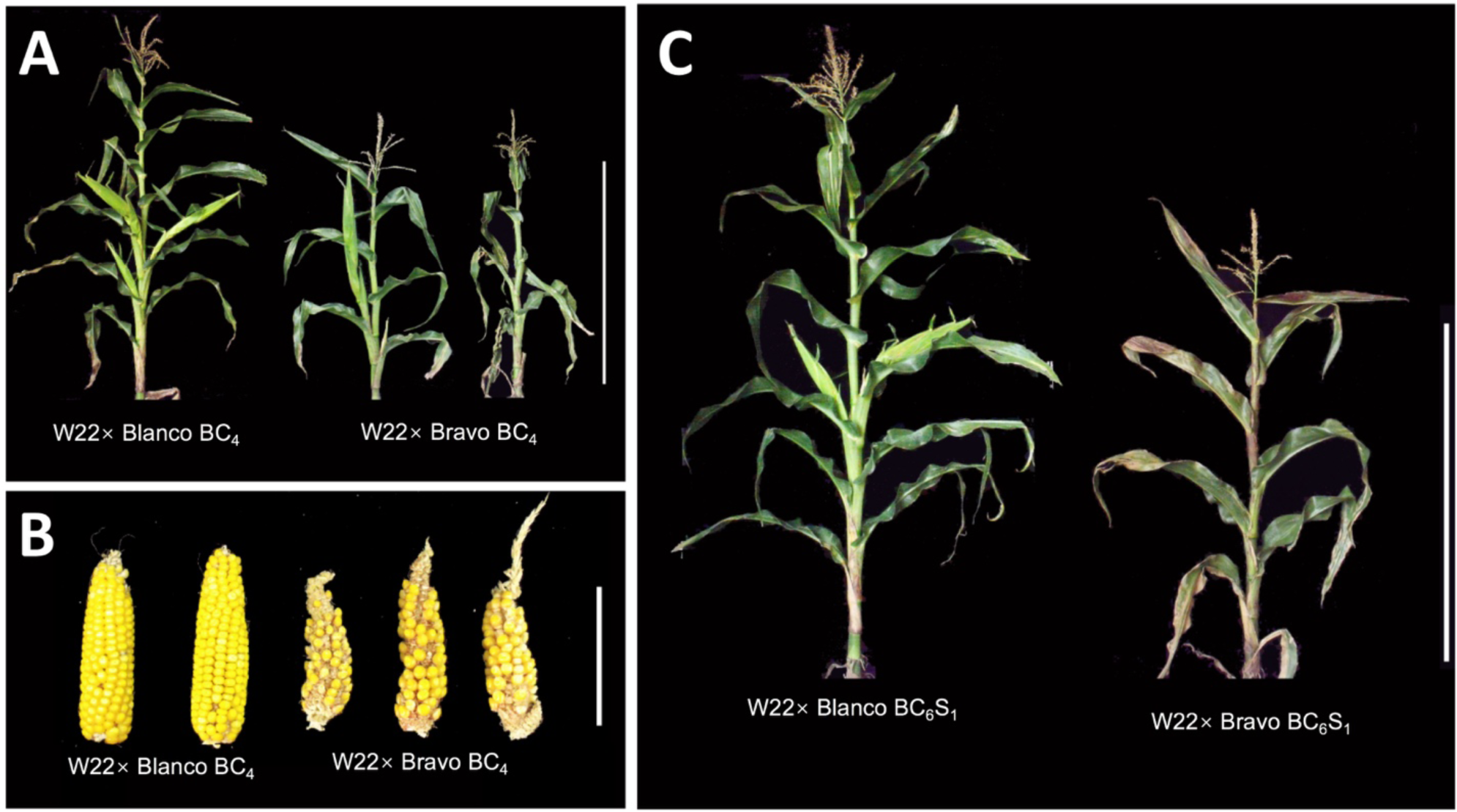
Phenotypes associated with the hybrid decay. Whole plants (A) and ears (B) from W22 x Bravo BC_4_ or a control accession W22 × Blanco BC_4_ are shown. In (C) examples of BC_6_S_1_ plants derived from Blanco or Bravo accessions are shown. Scale bars represent 1 m (A and C) and 10 cm (B).

Similar observations were made in an independent project attempting to generate BC_2_S_3_ recombinant inbred lines (RILs) from W22 and Bravo teosinte from the same F_1_ individual as used by Studer and Doebley (2012) (Figure 1A). In this project, roughly 2000 plants from the BC_1_ generation were grown (Briggs *et al.* 2007) and many of these BC_1_ plant showed aberrant phenotypes. The sickly syndrome became more apparent in the BC_2_ generation with all plants being clearly affected. Self-pollination of the BC_2_ plants yielded only sickly plants with no healthy segregants observed.

An independent set of crosses was performed to generate BC_4_ near-isogenic lines (NILs) between *Zea mays* ssp. *parviglumis* and maize inbred B73 (Liu *et al.* 2016). One of eleven accessions used for this study exhibited sickly phenotypes similar to that described above (Figure 1B). The sickly NIL population was derived from a teosinte accession (PI 384063; Beadle and Kato Site 7) collected 41 km south of Valle de Bravo (Figure S1), just 38 km from the location of the accession discussed above. In the lineages that exhibit the sickly syndrome, all plants were affected and there was no evidence for the syndrome to segregate. Difficulty in generating a sufficient number of progenies prevented advancing the sickly population beyond the BC_3_ generation, while the other ten NIL populations were created without problems.

In summary, these three experiments indicated that there is a previously unobserved phenomenon – hybrid decay – that appears in Bravo teosinte backcrosses to maize that causes a sickly syndrome that is transmitted from affected plants to all offspring across all subsequent generations of backcrossing and selfing.

### Confirmation and quantification of the hybrid decay syndrome

The independent discovery by two of our groups of hybrid decay with teosinte from the Valle de Bravo offered only anecdotal evidence. Therefore, we decided to confirm the occurrence of hybrid decay in crosses of maize and individuals of Bravo teosinte and compare these crosses to control crosses between maize and individuals from another teosinte population. For the control teosinte, we used an accession collected from 1 mile south of the town of Palo Blanco, Guerrero by Beadle and Kato (Site 4), hereafter “Blanco” teosinte. Blanco teosinte had previously exhibited normal phenotypic outcomes in crosses with W22 maize (Studer and Doebley 2012). We collected quantitative data on several phenotypes as a measure of the sickly syndrome.

First, we assessed whether the sickly syndrome could be documented in the BC_1_ of crosses of one of the Bravo accessions (Beadle & Kato Site 6) as well as Blanco teosinte (Beadle & Kato Site 4). The BC_1_ generation derived from the Bravo teosinte are shorter and have lower heights for the top ear relative to BC_1_ plants derived from the Blanco teosinte (Figure S3). Next, in order to provide a detailed description of the sickly syndrome while minimizing the effects of the segregation of genetic loci from maize and teosinte, we assessed a number of quantitative traits in the advanced W22 × Bravo backcross lines (BC_6-_ 8) compared to advanced BC lines derived from the cross of W22 and Blanco teosinte. These data showed that Bravo BC_6_ has a number of phenotypic abnormalities as compared to Blanco BC_6_ plants (Figure 3). Plant height and the height of the first ear node are significantly reduced in the Bravo BC_6_ and BC_7_ plants (Figure 3A). Nearly 50% of the plants have barren lateral branches in the Bravo BC_6_ and BC_7_ generations while barrenness was never observed in W22 or the Blanco BC lines (Figure 3A). The ears of the Bravo BC_6_ and BC_7_ plants have reduced diameter, length, kernel row number and seed set relative to the Blanco and W22 controls (Figure 3B). The seed set is nearly 3-fold lower for the Bravo BC materials. The Bravo plants have reduced tassel branch number, some male sterility and flower later than the controls (Figure S4A). The Bravo BC_8_ seeds also have reduced seedling root length at two days after germination as compared to Blanco BC_8_ seeds and W22 (Figure S4B).

**Figure 3.**
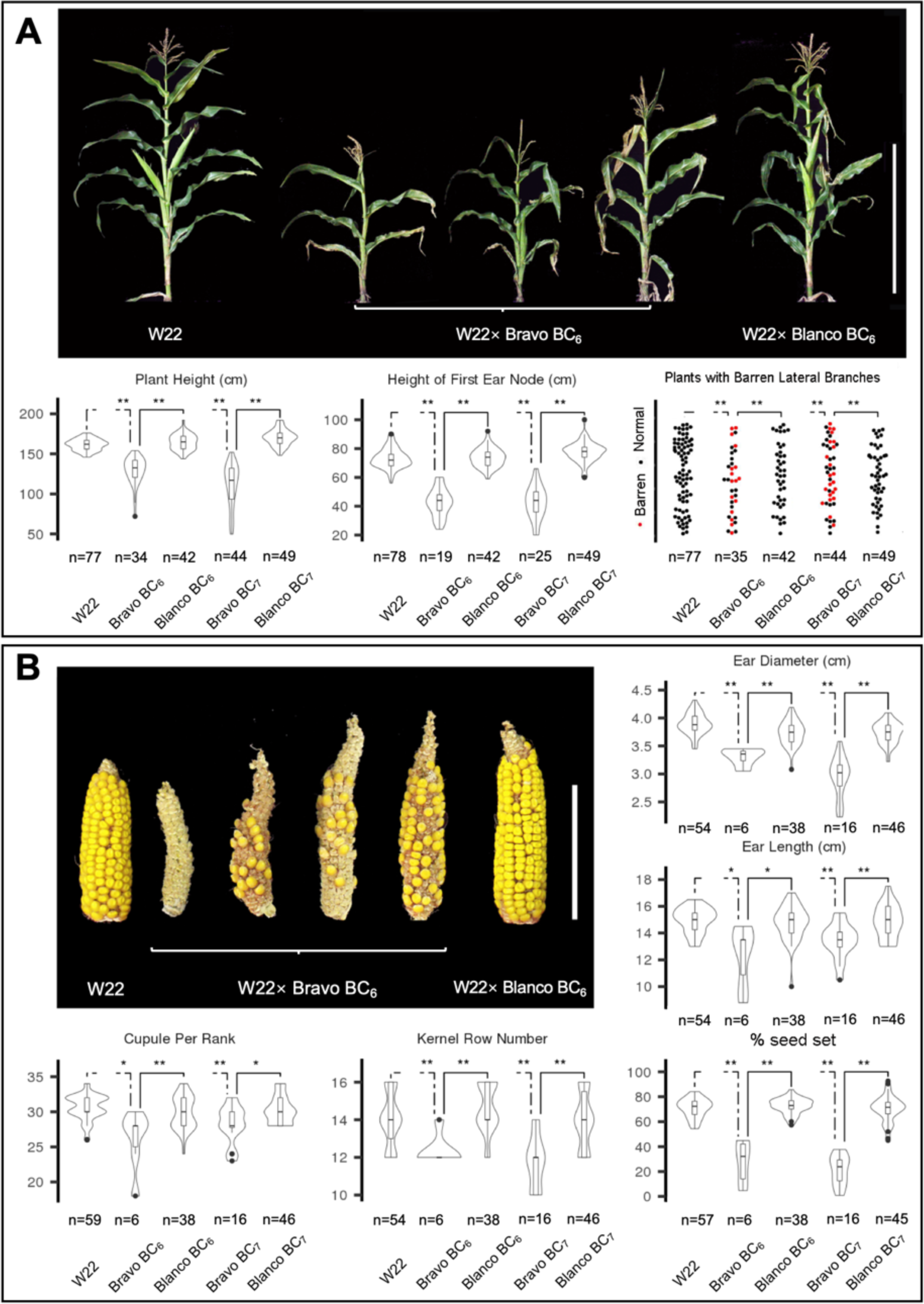
Phenotypic characterization of hybrid decay. (A) Plant morphology: examples of W22, W22 × Bravo BC6, and W22 × Blanco BC plants with violin plots for plant height, height of first ear node and proportion of plants with barren lateral branches. (B) Ear traits: ear diameter, ear length, cupules per rank, kernel row number and seed set with representative ears. Student’s t-test was used for plant height, the height of top ear node, ear diameter, ear length, and seed setting rate. Mann-Whitney-Wilcoxon test was used for cupule per rank, and kernel row number. *P < 0.05 and **P < 0.01. The number of plants measured for each trait (n) is listed for each plot. The two-dashed lines indicate comparisons between W22 × Bravo BC lines and W22 while solid lines indicate comparisons between W22 × Bravo BC lines and W22 × Blanco BC lines.

### Meiotic drive cannot explain hybrid decay

Meiotic drive (Lindholm *et al.* 2016), the preferential transmission of one allele from a heterozygote to all offspring, is a possible mechanism for hybrid decay. Under this mechanism, a sickly-syndrome-inducing allele would be transmitted to all offspring. A meiotic drive-based mechanism for hybrid decay would predict that a specific sequence from Bravo was retained through all BCs and was causal for the sickly syndrome. To test this hypothesis, we performed genotype-by-sequencing (GBS) for several individuals that exhibit the sickly syndrome as well as control plants (Figure S5A). Seeds from several different Bravo BC_6_ ears (3763, 3771 and 3772) that represent different backcrosses from a common Bravo BC_4_ line (Figure S5B) were screened. The BC_6_ plants tend to have one or two teosinte introgression regions per line, but we did not observe any specific teosinte introgression segments common to all Bravo BC_6_ plants as predicted by a meiotic drive mechanism (Figure S5A). This result suggests the sickly syndrome is not due to preferential inheritance of a specific locus through a meiotic drive like mechanism.

### Hybrid decay is not associated with karyotypic alterations

To test if there are any cytological manifestation of hybrid decay, we produced a FISH karyotype of the Blanco and Bravo teosintes, their W22 BC_6_ descendants and the recurrent W22 parent (Figure S6). The two teosinte karyotypes are distinct from each other (Albert *et al.* 2010) and are heterozygous for some sites. The distinctiveness of the teosinte karyotypes is expected. Both BC_6_ Bravo-derived plants have chromosomes typical of the W22 parent (Figure S6). The Bravo and Blanco BC_6_ progenies are comparable and show no evidence of chromosomal aberrations or abnormalities that distinguish the Bravo BC_6_ from the Blanco BC_6_ or W22.

### Hybrid decay is not dependent on transmission through male and female parents

The crossing schemes that uncovered hybrid decay were based on backcrossing using the recurrent maize line as the female (pistil) parent and the sickly plants as the male (pollen) parent. The mechanism of hybrid decay could potentially depend upon paternal transmission of genetic factors from sickly plants via their pollen. To test whether the sickly syndrome could be transmitted through both the male and female parents, the Bravo and Blanco BC_6_ lines were reciprocally crossed with W22. Plant height and height of the top ear node were equally reduced in the lines when the sickly Bravo BC_6_ line was used as either the male or female parent (Figure S7). This result suggests that once hybrid decay has been established, the sickly phenotype is not dependent on transmission through either the male or female parent. However, the direction of the original cross to create the F_1_ with maize as the female parent could be required to initiate hybrid decay.

### Genome changes in Bravo BC plants

We hypothesized that crossing Bravo teosinte and maize triggered genome instability such as the activation of transposable elements. In order to assess changes in copy number for genomic sequences in the Bravo BC plants, we assessed the Whole Genome Sequence (WGS) read depth for 1kb windows using low coverage WGS data aligned to the W22 genome (Table S1). The average read depth of the Bravo BC_1_ and Bravo BC_6_ plants relative to W22 (Figure 4A) reveals a handful of genomic regions with substantial (>10 fold) increases in copy number in the Bravo BC plants. The higher read depth for these regions likely indicates copy number gains, but does not necessarily mean that there are additional copies at this location. These additional copies could be located elsewhere in the genome.

**Figure 4.**
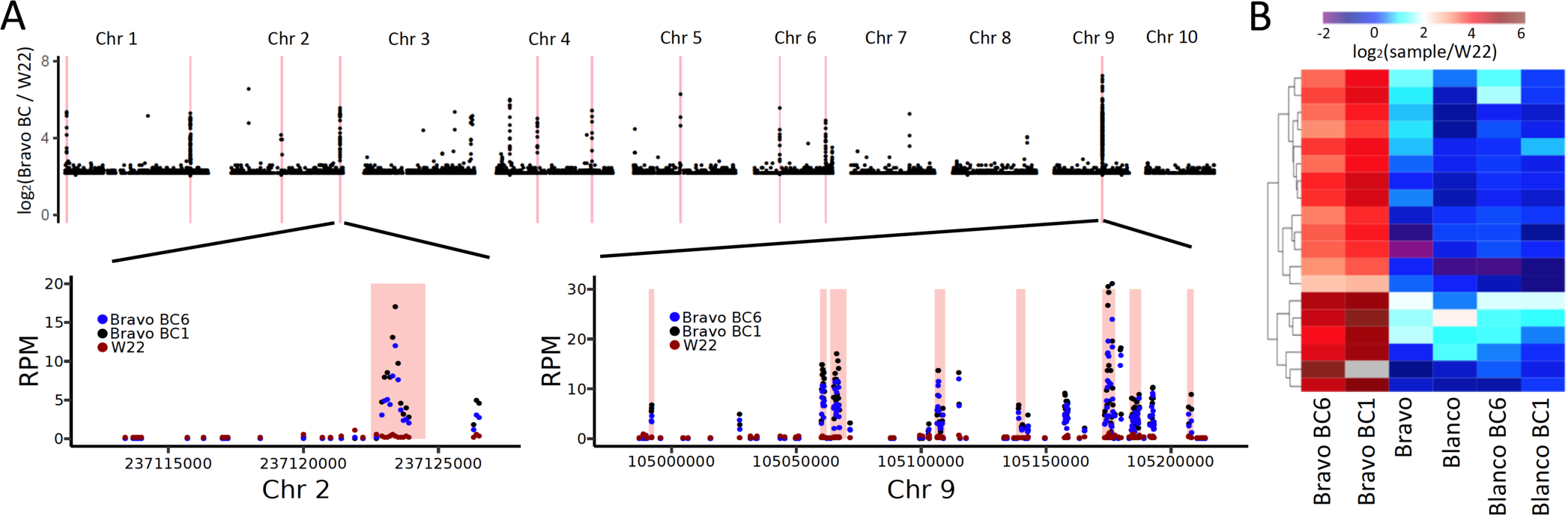
Genome content changes associated with the hybrid decay. (A) The read depth ratio (log2) for the Bravo BC (average of BC_1_ and BC_6_) relative to W22 is plotted for 1kbp bins across the genome. Only bins with differences >4-fold difference are shown. The red lines indicate regions that are found to exhibit consistently higher read depth in the Bravo BC materials relative to W22, Blanco and Blanco BC lines based on CNVSeq (P<0.01). For a region on chromosome 2 and a cluster of significant regions on chromosome 9, we show the reads per million (RPM) values for all windows with data in these regions. The red shaded regions indicate the regions identified as significant based on CNVSeq and the color of the points indicates the genotypes. (B) For each of these 19 regions, we determined the relative read depth of each sample relative to W22 and performed hierarchical clustering. The log2 (sample/W22) is indicated by the color. Dark blue indicates no change in read depth relative to W22 while red indicates higher read depth relative to W22.

To test which regions with aberrant read depth ratios in the Bravo BC plants relative to W22 were statistically significant, we used CNV-Seq (Xie and Tammi 2009). There are 19 regions that passed this test and all exhibit increased read depth in the Bravo BC lines relative to the other genotypes (Table S2; Figure 4). The regions that exhibit gains in read depth have 8-60-fold changes (Figure 4B) and are 1,000 - 6,500bp in length. The most prominent example is on chromosome 9 where there are eight significant CNV-Seq regions within a 50kb region (Table S2). The Bravo BC_6_ sample used for this analysis contains an introgression from Bravo in the middle of chromosome 7, but is otherwise homozygous for chromosomal segments derived from W22, therefore these are not likely to represent introgressed Bravo segments.

These 19 regions that exhibit consistently higher read depth in the Bravo BC samples also exhibit several notable features. First, they do not have high read depth in the sampled Bravo teosinte individual or W22 (Table S2; Figure 4B). This result suggests that these sequences were amplified in copy number during the backcrossing. Second, several of these sequences overlap annotated genes or transposons within the W22 genome. The boundaries of the regions showing copy number changes rarely align precisely with the boundaries of annotated features such as transposons or genes, however, limiting the ability to infer potential mechanisms for increased copy number. Third, visual inspection of the alignments in several of these regions suggests the higher copy is a sequence with homology but not identity to these regions (Figure S8). This inference is based on the observation that the elevation in read depth is variable within a single region and the observation that there are multiple SNPs relative to the W22 reference genome that are present in the majority of reads that are aligned to a region (Figure S8). In conclusion, our analysis of read mapping to the W22 genome identified sequences with similarity to some regions of the W22 genome that were increased in copy number during the backcrossing.

### Gene expression changes in Bravo BC plants

To identify changes in transcript abundance in sickly plants, we monitored steady-state transcript abundance using RNAseq in seedling tissue of individuals from Bravo-derived sickly plants and controls for both the W22 and B73 backcross series (Table S1). For the W22 × Bravo series, the W22 × Bravo BCs, the W22 × Blanco BCs (controls) as well as the Bravo, Blanco and W22 parents were assayed. For the B73 × Bravo series, we used a teosinte control from near the town of Teloloapan, Guerrero, Mexico (Accession PI384065) for which the BCs with B73 did not show the sickly syndrome (Liu *et al.* 2016). For this series, we assayed the B73 × Bravo BCs, the B73 × Teloloapan BCs (controls) as well as the Bravo, Teloloapan and B73 parents.

To understand the impact of hybrid decay on the transcriptome, we performed two analyses. First, a PCA (Principal Component Analysis) of the RNAseq data for all genotypes revealed that the plants fall into three clusters: W22 and its BCs, B73 and its BCs, and all pure teosintes (Figure S9A). Thus, the lines with the sickly syndrome cluster with their recurrent parent related lines, suggesting that this syndrome does not induce major differences in the W22 or B73 transcriptomes at the seedling stage. Second, to search for genes that are differentially expressed in sickly plants, DESeq (Love *et al.* 2014) was used to contrast the six sickly samples (W22 Bravo BC_1_ and BC_6_ and B73 Bravo BC_1_ and BC_2_) with the 14 non-sickly samples including the W22, B73, Bravo, Blanco, Teloloapan and control BC plants. This analysis identified a total of 18 DE genes (log2FC >1 and FDR<0.05), all of which are up-regulated in the Bravo BC sickly plants (Table S3, Figure S10). Thus, while the transcriptomes of plants with hybrid decay are not radically altered, there is a select set of 18 genes that are upregulated in these plants.

The 18 genes identified have some notable features. First, they do not appear to be randomly located around the genome but rather there is a cluster of five genes within a single 1 Mbp region on Chromosome 9 (Table S3). Second, Gene Ontology analysis does not show a specific functional category to which the 18 belong; indeed, ten of the 18 are hypothetical proteins with no known function. Third, only 8 of these 18 W22 genes have a collinear homolog annotated in the B73 genome and only 5 of the 18 genes have a collinear homolog based on comparison with sorghum (Table S3). Genes that lack a collinear homolog in other maize lines (and sorghum or other grasses) are typically non-essential. There are relatively few examples of mutants in such genes that have a detectable phenotypic effect (Schnable 2015). Thus, although these 18 genes cannot be excluded as candidates for the causal agents of hybrid decay, their sequence homology provides no insight into how they might cause it.

The RNAseq alignment data for each locus revealed a few key details (see examples in Figure S11). First, only four of the 18 genes exhibit read mapping patterns suggestive of expected transcription throughout the gene and proper splicing. Second, for 14 of the 18 genes, the increase in transcript abundance does not match the predicted structure of the W22 transcript, rather they exhibit expression only for a portion of the gene, often overlapping an intron (Figure S11). Third, the majority of reads that map to these 18 genes have sequence polymorphisms relative to the W22 genome. This observation suggests that the increased expression is likely from sequences from the Bravo parent with homology to several regions of W22 but is not likely due to transcription of sequences that are present in W22.

### Identification of novel transcripts present within Bravo BC plants

The observation that some of the 18 W22 genes with increased expression were similar to, but distinct from, W22 genome sequences led us to investigate the possibility of novel transcripts in these plants. We performed a *de novo* assembly of transcripts from the RNAseq data for sickly plants. The reads from the W22 Bravo BC_1_ and BC_6_ samples were pooled and used for transcript assembly by Trinity (Haas *et al.* 2013). We obtained 73,145 assembled transcript contigs (with 93,592 isoforms) from this assembly with an N50 of 1,358bp. Then, a PCA using the read counts per transcript was performed and revealed that PC2 separates the sickly plants from other related individuals (Figure S9B). The ability to distinguish sickly from normal plants with a PCA using these novel transcripts implies that the sickly phenotype involves transcripts that are not part of the W22 gene set. Next, to identify specific sequences that are differentially expressed in the sickly plants relative to the other genotypes, all RNAseq reads from all samples were aligned to assembled transcripts. Differential expression analysis identified 59 transcripts that have significant differences in expression between the 6 sickly samples relative to the 14 non-sickly samples (Table S4; Figure 5).

**Figure 5.**
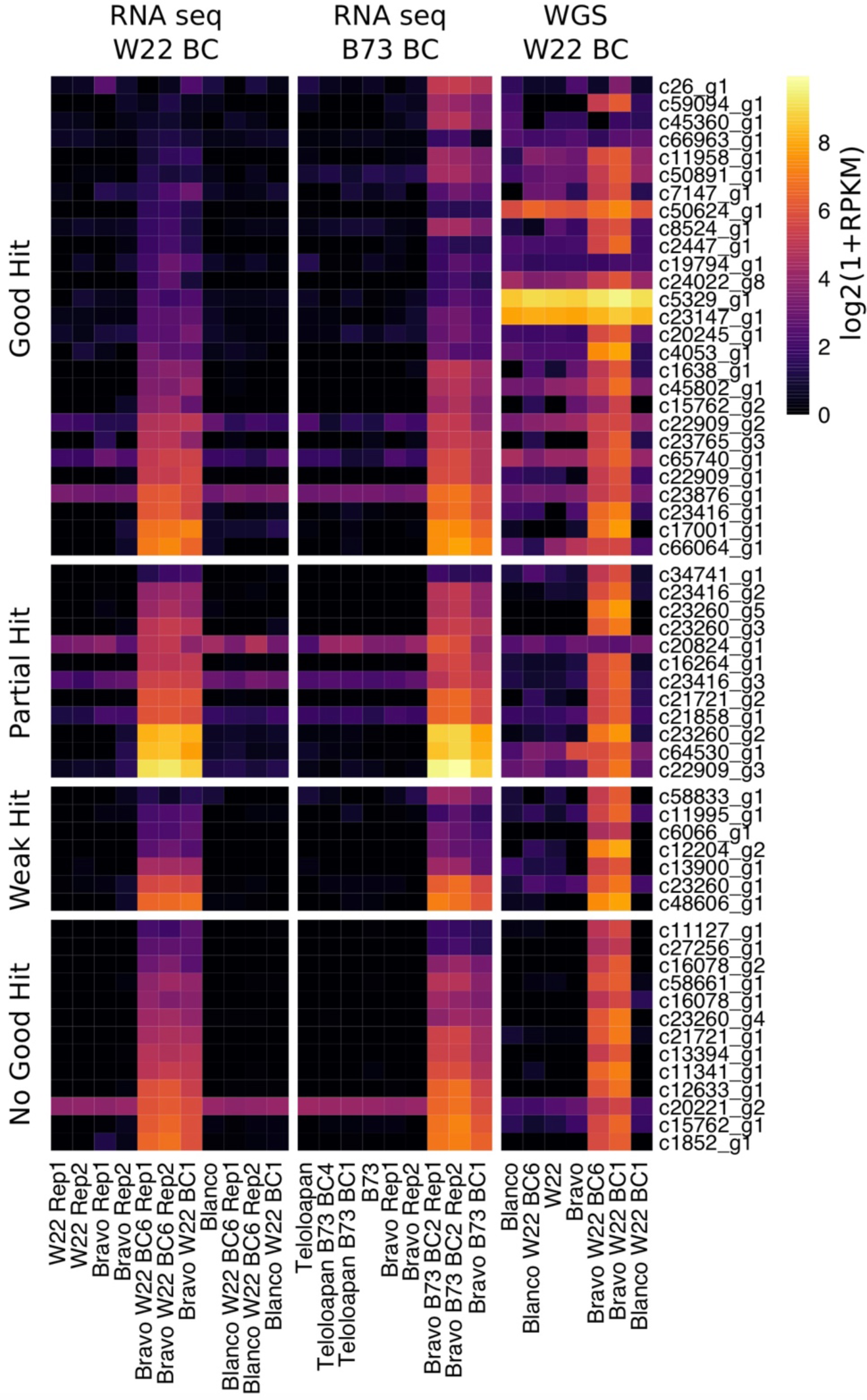
Expression and read depth for *de novo* transcripts with altered expression in sickly plants. The *de novo* assembled transcripts (from W22 Bravo BC_1_ and BC_6_ RNAseq samples) were used to perform differential expression analysis. A set of 59 transcripts with altered expression was identified. These transcripts were classified based on their alignments to the W22 genome. Each group of transcripts is ordered by FPKM expression in the Bravo BC_6_ Rep1 sample and a heat map is used to visualize the expression level. In addition, the RPKM values based on unique alignments of the WGS data to all *de novo* transcripts are also visualized as a heatmap using the legend on the right.

We proceeded to characterize these 59 transcripts that are highly expressed in seedling tissue of the sickly genotypes (Table S4). In order to determine whether these transcripts represent genes that are present in the W22 genome, genes that are highly similar to W22, or novel sequences, the highest expressed isoform of each transcript was aligned to the W22 genome using BLAST. There are 27 transcripts with high similarity to the W22 genome (>95% identity and >50% of the length of the transcript aligning to the genome). There are 12 transcripts with partial alignments to the W22 genome (>95% identity but less than 50% of the transcript aligns to the genome). The remaining 20 transcripts have either weak BLAST hits (7 transcripts with <95% identity) or no significant alignments to the W22 genome (13 transcripts). The annotations for the genomic regions that the 39 transcripts with good or partial hits to W22 were assessed. There are 10 transcripts that at least partially overlap annotated genes (including 8 of the 18 W22 genes we identified). The remaining 29 transcripts included 16 that overlap annotated retrotransposons (RL), 3 that overlap helitron (DH) elements, and 10 that align to unannotated regions between genes and TEs. In many instances, these transcripts only overlap a portion of these features (Table S4) and likely do not represent transcription of the full annotated gene or transposable element although there are some examples of the transcript containing the full features.

There are questions about the potential functions that might be encoded in these up-regulated transcripts. The coding potential and domains present within these transcripts were assessed. Only about half of the highly expressed transcripts have coding potential and 13 of these produce putative proteins that contain domains with significant similarity to PFAM domains (Table S4). There was little evidence for a common function that was present in these transcripts. There are examples of specific enzymatic activities (amino transferase, glycosyl transferase, peptidase) or domains of unknown function. One of the more interesting observations was the presence of several long transcripts (c23147_g1_i2 and c23260_g1_i7) that contain regions annotated as having retrotransposon functions (integrase/reverse transcriptase) and another transcript (c13394_g1_i3) with a transposase related domain. The observations that several of the transcripts contain domains related to transposable elements and that some transcripts align to regions annotated as transposable elements suggest there is up-regulation and activation of transposable elements in hybrid decay.

### Correspondence between upregulated genes/transcripts and WGS CNVs

Since the RNAseq expression data for both the 18 W22 genes and 59 novel transcripts suggested that something more complicated than simple gene up-regulation was occurring and since the WGS data indicated that some regions of the genome had elevated copy number in sickly plants, we compared the RNAseq fold change (log_2_) values to that for the WGS data. The read depth of each of these genes/transcripts was assessed by aligning the WGS data to the 18 W22 genes and the 59 *de novo* transcripts and determining FPKM values (Tables S3 and S4). First, Figure S10 shows that the 18 upregulated genes in sickly plants exhibit higher read depth in the whole-genome shotgun data, suggesting increased copy number at the DNA level. Second, the majority of 59 novel transcripts have a higher WGS read depth in sickly plants relative to non-sickly plants (Figure 5; Table S4). There are 41 transcripts with at least 10-fold increase in read depth in the Bravo BC plants and another 11 transcripts with 3-10-fold increases in read depth. It is notable that the majority of these regions exhibit relatively low copy in the Bravo teosinte individual that was sequenced (Figure 5). In conclusion, there is a set of genes/transcripts that are highly expressed and increased in copy number in the sickly plants and many of these sequences are not present in the W22 genome, suggesting a proliferation of sequences donated by the Bravo teosinte parent.

### Correspondence of up-regulated sequences to transposons

To investigate the potential up-regulation of transposable elements (TEs) from the W22 genome in the Bravo BC plants, we mapped the RNAseq reads to transposons using a mapping strategy that counts the number of RNAseq reads per transposon family rather than per locus (Anderson *et al.* 2018). To enable comparisons between the RNAseq and WGS datasets, we tested for a difference in counts per TE family between sickly Bravo BCs (BC_1_ and BC_6_) and normal Blanco BCs (BC_1_ and BC_6_) (Tables S5 and S6). This analysis identified 11 TE families with significant increases in expression (Table S5) and 18 TE families with increase in read depth for WGS data (Table S6). Seven of the TE families were identified as having significant increases in both expression and genomic read depth. A comparison of the expression levels and genomic read depth for each of these families in all samples reveals that the increases in expression and/or read depth are limited to the Bravo BC families (Figure 6). Unclassified LTR retrotransposons (RLX) were the most common families that exhibited increase. However, there was also a Helitron family (DHH) and two gypsy-like retrotransposons families (RLG) with increases in both expression and copy number (Tables S5 and S6). There are no TE families with an elevation of expression or genomic copy number in the normal Blanco BCs. Thus, at least 22 TE families show evidence of activation in sickly plants.

**Figure 6.**
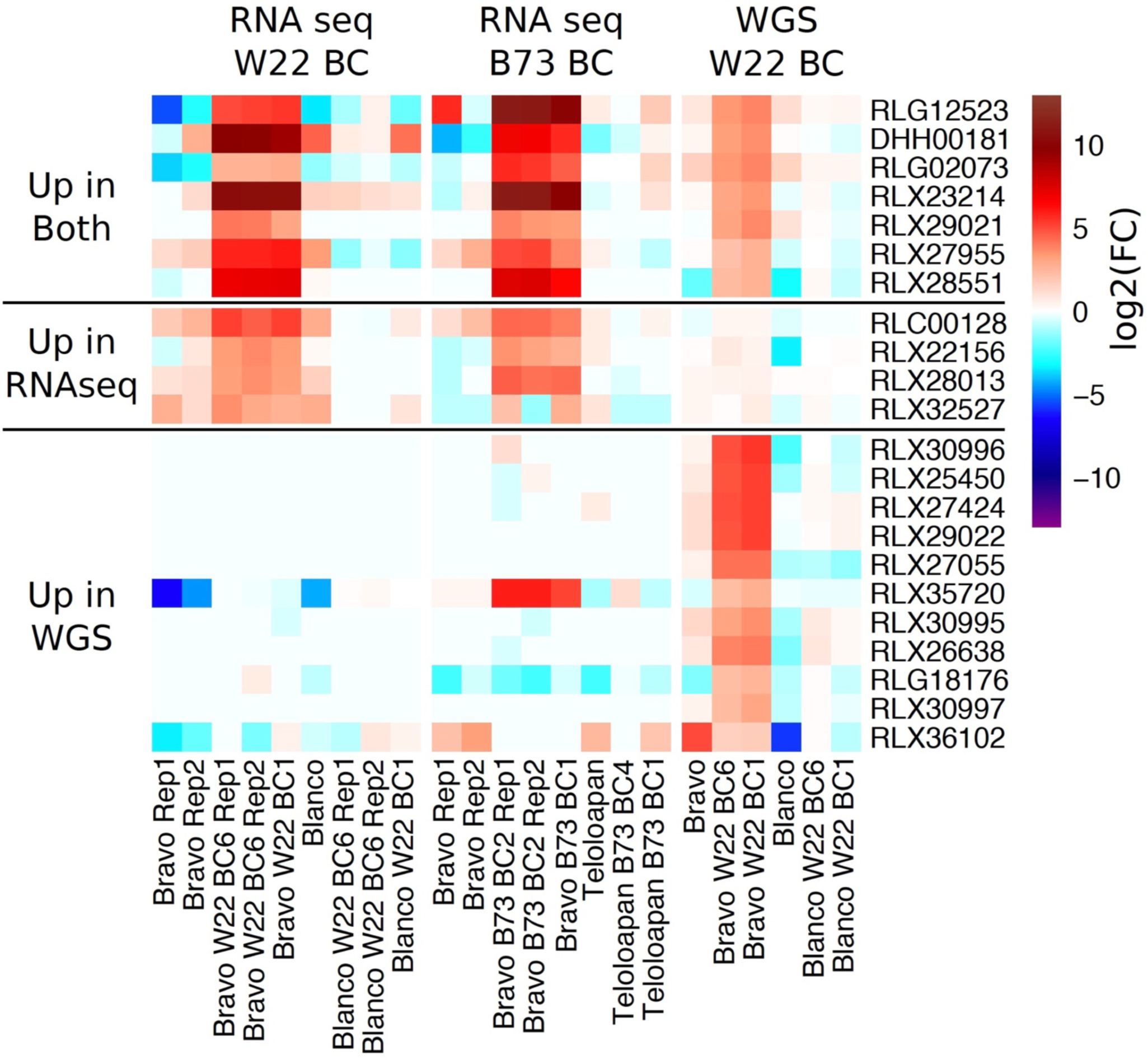
Changes in expression and copy number for some transposable element (TE) families. TE families that exhibit increases in expression level or read depth were identified by comparing the W22 Bravo BC_1_/BC_6_ lines with W22 Blanco BC_1_/BC_6_ controls. A clustered heat map is used to visualize expression and copy number relative to W22 for these TE families in all samples. The TE families were divided into three groups, the 7 TE families that exhibit significant increase in both RNAseq and WGS data, four families with significant increase in RNAseq but not in WGS data, and the 11 families with significant increase in WGS data but not in RNAseq.

### DNA methylation changes in plants with the hybrid decay syndrome

In order to test for changes in DNA methylation in sickly plants, we generated whole genome bisulfite sequencing (WGBS) data for the W22 BC materials to search for unique DNA methylation patterns in sickly plants. The analysis of DNA methylation patterns was restricted to alignments of reads to the W22 genome. This approach provides a survey for the majority of genome present in the W22 BC samples. A comparison of the global DNA methylation patterns by PCA reveals that the Bravo BC plants are closely related to W22 with no evidence for genome-wide repatterning of DNA methylation (Figure S12). To identify loci with altered DNA methylation, the methylome of the Bravo BC_1_ and BC_6_ plants was compared to the W22, Bravo teosinte and Blanco BC_6_ plants to identify CG and CHG differentially methylated regions (DMRs) with >60% difference in DNA methylation levels for 100bp tiles. This analysis identified 167 CG DMR regions (Table S7) and 260 CHG DMR regions (Table S8). Reduced methylation accounted for 75.4% of CG DMRs and 62.3% of CHG DMRs. There are CG DMRs near 11 of the 19 regions with elevated copy number. There are also CG DMRs located within 2kb for 19 of the 46 up-regulated transcripts that had alignments to the W22 genome (Table S4).

### sRNA abundance in plants with the hybrid decay syndrome

We analyzed the genome-wide changes of sRNAs using small RNA-sequencing (sRNA-seq) at the seedling stage of the sickly Bravo BC plants and normal control plants for both W22 and B73 background (Table S1). We did not observe global changes of sRNAs between normal and sickly plants in either genetic background (Figure S13-S14), suggesting that the sickly phenotype is not associated with major disruption of sRNAs. To identify genomic regions with differential levels of sRNA abundance between normal and sickly plants, we quantified sRNA expression by normalizing against total reads for each size class of sRNAs per 500-bp window along the W22 and B73 genomes. A total of 200 and 112 regions (FDR ≤ 0.01 & fold change ≥ 2) with differential sRNA abundance were identified in the W22 and B73 genomes, respectively (Table S9-S10). Most of the differentially expressed sRNA regions showed strongly directional bias to significantly increased sRNA levels in sickly Bravo BC plants in both W22 and B73 background, especially for 21 and 24-nt sRNAs (Figures 7A, S15A; Table S9-S10). More than half of these regions exhibit up-regulation for multiple sizes of sRNAs simultaneously in sickly plants as compared to normal control plants (Figure 7A, B, S15A, B). Although we also observed that some of regions in sickly maize with W22 genetic background specifically produced decreased 22-nt sRNA compared with the normal plants, this is not the case for the B73 genetic background (Figure 7A, C and S15A). Thus, we concluded that a group of sRNAs produced in the seedlings that will later exhibit a sickly phenotype might contribute to hybrid decay in Bravo BC plants.

**Figure 7.**
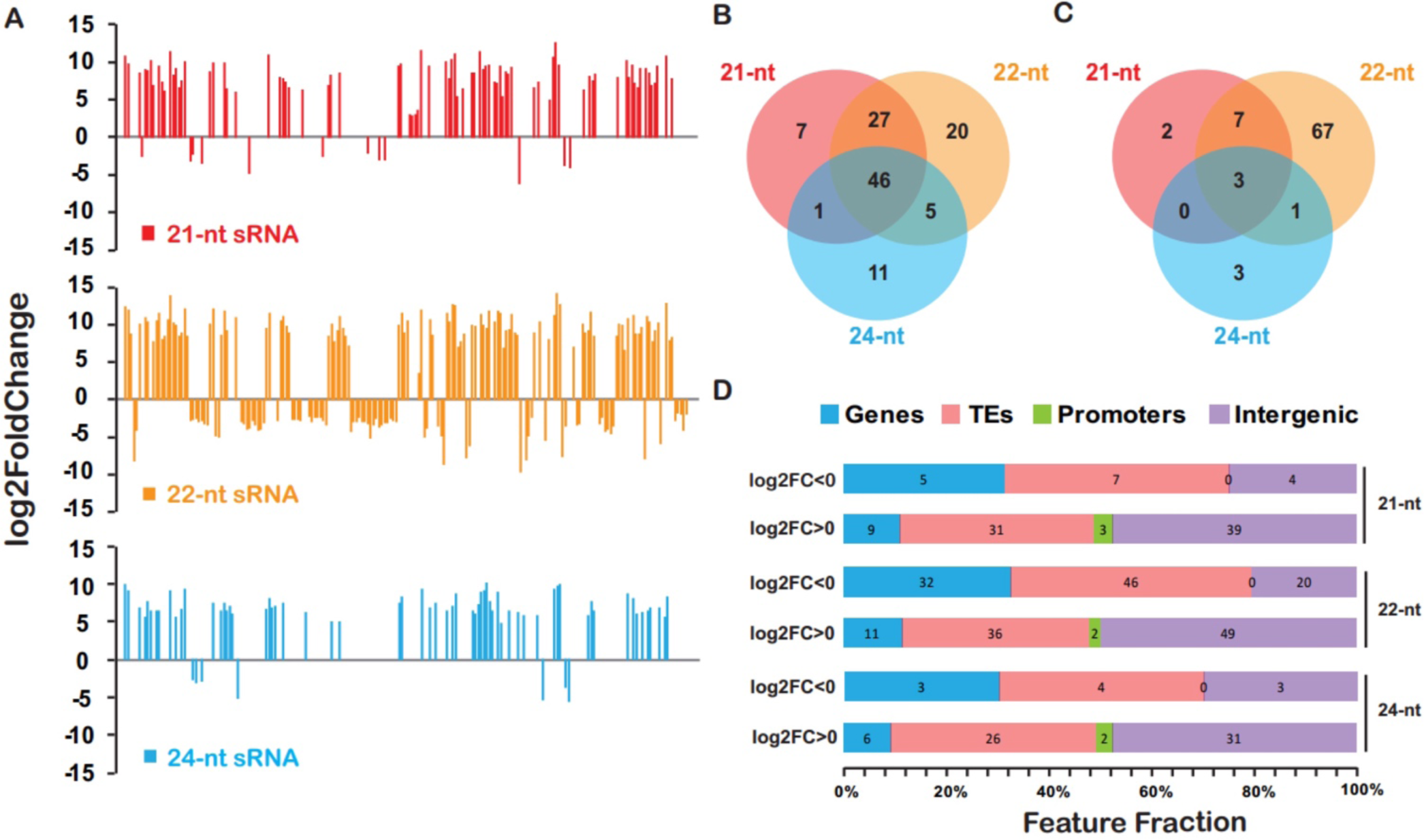
Feature analyses of differentially expressed sRNA regions in sickly Bravo BC plants compared with controls. (A) Graph of differentially expressed sRNA regions one-by-one, showing the bias that most of differentially expressed sRNA regions produced increased 21-nt, 22-nt, and 24-nt sRNA in sickly plants as compared to control plants. The y-axis indicates the log2 transformation of fold changes, with positive value representing the generation of increased sRNA in sickly Bravo BC plants compared with the normal controls. (B) Almost half of differentially expressed sRNA regions with increased sRNA in sickly plants produced three classes (21-nt, 22-nt, and 24-nt) of sRNAs concurrently. (C) Most of differentially expressed sRNA regions with decreased sRNA in sickly plants only produced 22-nt sRNAs. (D) Overlap profiling of differentially expressed sRNA regions with W22 genomic features, including genes, transposable elements (TEs), 2-kb promoter regions of annotated genes, and intergenic regions. The log2FC >0 means that increased sRNAs were generated in sickly Bravo BC plants compared with the normal controls. The log2FC <0 means that reduced sRNAs were produced in sickly Bravo BC plants compared with the normal controls.

An analysis of the differential sRNA generating loci showed overlap with genomic features, such as transposable elements (TEs), gene body and promoter regions of annotated W22 or B73 genes (Figure 7D, S15C). Thus, we re-analyzed the sRNA changes with TEs or genes as units of measurement (Tables S11-S14). A total of 25 annotated W22 genes showed significantly differential accumulation of sRNAs between normal and sickly plants (Figure 8A; Table S11). Similarly, we detected 13 annotated B73 genes with a different amount of sRNAs generated between normal and sickly plants (Figure S16; Table S13). Most of these genes exhibit significant variation for 22-nt sRNA levels (Figure 8A). Only five genes with colinear homology show altered sRNA profiles in both W22 and B73 - four that produced increased and one that produced decreased sRNAs in sickly plants (Figure 8A, 15A; Table S11, S13). We also detected some TEs with altered sRNA generation in sickly plants compared with normal plants (Figure 8B; Table S12, S14). Interestingly, almost all of these TEs consistently produced more sRNA in sickly plants than in normal plants for both W22 and B73 background (Figure 8B, S15B). Strikingly, four of these TEs also showed either increased CNV, up-regulated expression of novel transcripts, and reduced CG and/or CHG DNA methylation around them (Figure 8B). We speculate that hybrid decay affects DNA methylation around these TEs so that they and neighboring regions express novel transcripts or produce extra sRNAs.

**Figure 8.**
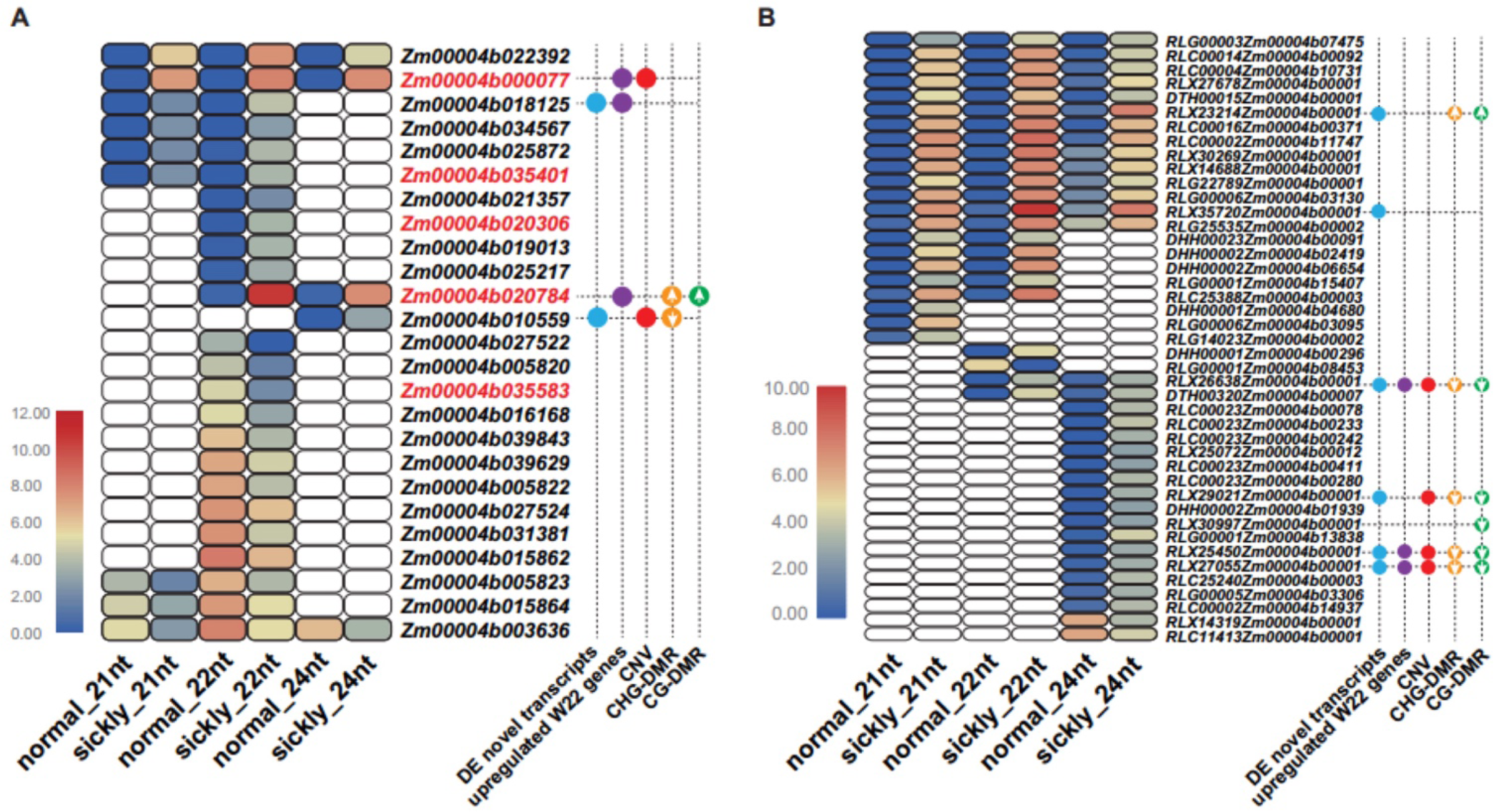
Annotated W22 genes (A) and transposable elements (TEs) (B) that potentially generated significantly different amount of sRNAs between sickly and control plants and their comparisons with other features of genes or novel transcripts expression, CG or CHG DNA methylations, and copy number. Genes with red fonts indicate that the corresponding B73 homolog genes also produced more sRNA in sickly plants with B73 genetic background (Figure S16). The points with different colors stand for genes or TEs that are located near the novel transcripts (blue), up-regulated W22 genes (purple), increased copy number regions (red), and increased or decreased CG (orange) or CHG (green) methylation regions in sickly plants with W22 background.

## Discussion

Hybrid decay is a transgenerational epigenetic phenomenon observed in the backcross progeny of certain teosinte individuals from near the Valle de Bravo with W22 or B73. Abnormal, sickly phenotypes appear in the BC_1_ generation and a pronounced sickly phenotype is manifest in more advanced BC generations. The phenotypic effects of hybrid decay are pleiotropic, affecting plant stature, the ear, the tassel and root growth. A remarkable feature of hybrid decay is that the Bravo and maize parent lines are themselves normal phenotypically, indicating that hybrid decay is due to an interaction between these two genomes and not a feature of either one. We also observed that this syndrome once established can be transmitted to offspring through either the male or female gametes. The inheritance of the hybrid decay is non-Mendelian and epigenetic in that it is transmitted from parent to all progeny and does not segregate among the progeny. Once the syndrome is initiated in a lineage, normal plants are never recovered in any progeny of subsequent generations.

Hybrid decay alters the genome of affected plants in multiple ways. First, there are some genomic sequences with homology to 19 regions of the W22 genome that have strongly elevated copy numbers in sickly plants. Eight of these sequences are clustered in a 50 kb segment of chromosome 9. Second, these amplified sequences do not have exact sequence identity to the W22 genome but show sequence polymorphisms that distinguish them from W22, suggesting that their origin may be from the Bravo genome. Third, there are 18 W22 genes that are up-regulated in plants with the sickly syndrome, and many of these genes share sequence homology with the amplified genomic sequences and some also map to the same region on chromosome 9. Fourth, *s*ickly plants express 59 novel transcripts, all of which are up-regulated in sickly plants as compared to controls. Some of these transcripts share homology with the 18 differentially expressed W22 genes or the genomic sequences that have elevated copy number in sickly plants. Fifth, there are 22 TE families that have either elevated expression, elevated WGS read depth or both. Most of these are retrotransposons. Sixth, there are 231 differentially methylated regions between sickly plants and heathy controls. Many of these regions, which show reduced methylation in sickly plants, are also associated elevated copy number in sickly plants. Seventh, there are several hundred genomic regions with differential sRNA production, most of these have elevated levels of sRNAs in sickly plants. Finally, these multiple genomic features of hybrid decay are correlated such that some W22 genes and TEs involved show multiple anomalous features.

A key feature of hybrid decay is that after 6 generations of backcrossing to maize, it is expected that the genome of Bravo-derived plants will be largely identical to that of the recurrent parent with only small segments of residual Bravo DNA as regions of heterozygosity. Both FISH and GBS marker data confirm that this is true. However, we also detected novel transcripts in the sickly BC_6_ plants that are not present in the recurrent parent. CNVs in the sickly plants demonstrate that some genomic sequences are being amplified. While these CNVs are similar to sequences present in the W22 genome, they are clearly not derived from the W22 genome based on the presence of polymorphisms. Moreover, many of the novel transcripts have low homology to the W22 genome. Together, these observations suggest that non-W22 sequences have proliferated in the genome of the Bravo BC plants, that these sequences can be highly expressed and associated with alterations in sRNA production and DNA methylation. Some of these sequences have homology to transposable elements and it is possible that transposon activation is involved in this genomic instability.

### Some unanswered questions

A central question for future research will be to identify the sequence(s) in Bravo teosinte and/or maize that act as the driver(s) that triggers hybrid decay. If there are specific sequences or DNA elements that trigger hybrid decay, why do they not initiate this phenomenon in the pure Bravo or the maize parent? Is there a protective factor (or epigenetic state) that silences the drivers in Bravo teosinte that is not transmitted from the F_1_ to the BC generations? Are the drivers activated in the F_1_ such that its gametes are set to trigger hybrid decay or is this phenomenon initiated in the zygote of the BC_1_? Would a Bravo teosinte × W22 F_2_ population exhibit hybrid decay or even segregate for it? What are the steps involved from the initiation of hybrid decay to the likely downstream effects that we describe including elevated expression of novel transcripts, CNVs, amplification of transposable elements and changes in the methylation and sRNA profiles? We observed a significant concentration of genomic alteration associated with a region on chromosome 9. Is there something special about this region regarding hybrid decay? Regarding the CNVs, are they the result of tandem duplication or elements dispersed throughout the genome?

Other questions to be answered include: (1) Is Bravo teosinte polymorphic in this regard such that some Bravo teosinte individuals would not trigger hybrid decay? (2) Are there other teosinte populations that can trigger hybrid decay? (3) Our crosses were all made using maize as the female parent. Would hybrid decay also have been triggered if maize had been used as the pollen parent and teosinte as the female parent?

### Thoughts on the underlying mechanism

We suspect that the activation and amplification of TEs from the Bravo genome in the Bravo BC plants underlie hybrid decay and that these TEs might have escaped genome surveillance during hybridization. Many TEs in plant genomes are kept silent by DNA methylation at CG, CHG, and CHH contexts (Law and Jacobsen 2010; Matzke *et al.* 2015; Cuerda-Gil and Slotkin 2016). *De novo* methylation occurs through a mechanism known as RNA-directed DNA methylation (RdDM), in which siRNAs derived from TEs guide the deposition of DNA methylation at homologous sequences (Law and Jacobsen 2010; Matzke *et al.* 2015; Cuerda-Gil and Slotkin 2016). Once established, methylation at CG contexts can be maintained independently of RdDM, but methylation at CHG and particularly CHH contexts requires RdDM to be maintained (Law and Jacobsen 2010; Matzke *et al.* 2015; Cuerda-Gil and Slotkin 2016). *De novo* CHH methylation has been shown to be essential in preventing new bursts of transposition in *Arabidopsis* (Marí-Ordóñez et al., 2013; Cavrak et al., 2014). In this context, it may be significant that a CHH methyltransferase (Zm00004b000077 in W22; Zm00001d027329 in B73) is among the genes with elevated mRNA and sRNA levels in sickly plants (Figures 8A, S16A).

During plant development, DNA methylation is reprogramed and reinforced in the germ cells and during embryogenesis. The central cell but not the egg cell undergoes active demethylation, resulting in the activation of TEs and the production of siRNAs in the endosperm. The siRNAs are thought to move from the endosperm into the embryo to cause *de novo* DNA methylation at homologous TEs (Hsieh *et al.* 2009; Bouyer *et al.* 2017). Studies in *Arabidopsis* show that CHH methylation in embryos increases during embryogenesis, reaching full methylation levels in mature embryos (Bouyer *et al.* 2017; Kawakatsu *et al.* 2017). CHH methylation levels decline subsequently in plant development because the RdDM machinery is highly expressed only in meristematic tissues (Hsieh *et al.* 2009; Bouyer *et al.* 2017). Thus, embryogenesis is a key period when DNA methylation is reinforced. We speculate that, in the F_1_ embryos of the maize x Bravo cross, certain TEs in the Bravo genome cannot be properly targeted by maternal siRNAs derived from maize TEs in the endosperm due to sequence diversification. These TEs are thus amplified and inserted into the maize genome, as revealed by CNV-seq in this study. In post-embryonic F_1_ plants, although the TEs give rise to siRNAs, which are likely derived from RNA polymerase II-generated transcripts from the TEs, the siRNAs are unable to cause DNA methylation. Potential reasons include low levels of expression of RNA polymerase V (Pol V), which is required for siRNA-mediated DNA methylation, and inability of Pol V to access active chromatin at the TEs (note that DNA methylation promotes the recruitment of Pol V to chromatin in *Arabidopsis*). In subsequent BCs, the TEs continue to escape surveillance due to the lack of maternal, maize-derived siRNAs that can target them, resulting in their further amplification and remobilization.

How can TE activation and remobilization cause hybrid decay? Two possible mechanisms are proposed here. First, in our present study, we observed sRNAs of 21-24 nt that are specifically produced from the strongly amplified TEs or TE homologs in sickly plants (Figure 8). We speculate that these “foreign” or “ectopic” sRNAs may repress the expression of certain maize genes to cause the sickly phenotype. For example, the 24-nt siRNAs might target some maize sequences and cause DNA methylation, which in turn could repress the expression of genes near these sites. The 21-nt siRNAs may target maize transcripts to cause post-transcriptional gene silencing through RNA cleavage. The 22-nt siRNAs could even lead to the biogenesis of secondary siRNAs from target transcripts, and the secondary siRNAs may have additional targets in the genome, thus amplifying the effects. In summary, the down-regulation of target genes by the ectopic siRNAs leads to the sickly phenotype. Although down-regulation of gene expression was not observed with our RNA-seq data from seedling tissues, one cannot rule out that this occurs in specific tissues or cells that are crucial for development, such as meristems. Alternatively, the ectopic siRNAs could lead to translational repression of maize mRNAs, an effect that is not reflected at the transcriptome level. Second, the activated TEs could lead to the up-regulation of nearby genes to cause the sickly phenotype. It has been documented that the epigenetic states of TEs can affect the expression of nearby genes, especially when the TEs are in promoters or introns of genes (Ito and Kakutani 2014; Cui and Cao 2014). In this study, we observed up-regulation of 18 W22 genes as well as 59 novel transcripts that are probably from the Bravo genome. These protein products may have adverse effects on plant development. Additionally, there may be specific upregulated sequences that contribute to hybrid decay. We note that one LTR retrotransposon family (RLX23214) with increased copy number in sickly plants carries a sequence homologous to a CHH methyltransferase gene (Zm00004b000077 in W22; Zm00001d027329 in B73), and sRNAs map to regions of both the TE and the gene. Although knockouts of this gene do not have an obvious phenotypic effect in maize inbreds (Li et al., 2014), the gene is an active component of silencing in reproductive tissues (Garcia-Aguilar et al., 2010) and provides a compelling target for continued study of the mechanistic basis of hybrid decay.

Finally, we note that our observations of hybrid decay with teosinte are similar to a form of hybrid dysgenesis observed in maize. When Zapalote Chico, a maize landrace grown by the Zapotec people of Oaxaca, is crossed to other maize germplasm, offspring show low vigor or sterility (Gutierrez-Nava et al., 1998). This hybrid dysgenesis is due to the activation of a *Mutator* transposon in crosses with non-Zapalote Chico maize germplasm (Gutierrez-Nava et al., 1998). As in hybrid decay, the initial cause is unknown, but the outcome – differential TE activity and methylation – is similar. Understanding the epigenetic and genetic similarities of systems of hybrid decay and hybrid dysgenesis may help to understand the prevalence of these systems, and if they play a role in reproductive isolation and divergence between populations and species.

## Material and Methods

### Plant materials, ***RNA preparation***, ***and DNA preparation***

For backcrosses to W22, a Bravo teosinte accession (Beadle and Kato Site 6 from 79 km south of Valle de Bravo) was used, and a Blanco teosinte accession (Beadle and Kato Site 4 from 1 mile south of the town of Palo Blanco, Guerrero) was used at the normal control. For backcrosses to B73, another Bravo teosinte accession (PI384063; Beadle and Kato Site 7, 41 km south of Valle de Bravo) was used, and a Teloloapan teosinte accession (PI384065) was used as the normal control. The W22 backcrosses for both Bravo teosinte and the Blanco teosinte control were generated as part of a previously published project (Studer and Doebley 2012). The B73 backcrosses for both Bravo teosinte and the Teloloapan control were generated as part of another previously published project (Liu *et al.* 2016). The BC lines derived from both Bravo teosinte accessions were not included in these published papers because of the unexpected hybrid decay phenomenon, however they were constructed as part of those projects as were the BC lines for the teosinte controls. For the current project, the W22 BC_6_ lines were crossed with their recurrent parent to produce BC_7_ (and subsequent BC_8_) seed.

In order to sample plant tissues for genomic analyses, seeds were treated with fungicide and placed on multiple layers of damp germination paper and covered by one layer of wet germination paper. The tray was placed under fluorescent lights at the room temperature. When plants reached ∼5 cm height, they were transferred to small pots containing soil in the growth chamber with an 11-hour light and 13-hour dark cycle until the V1 to V2 seedling stage. Then whole above-ground seedlings were harvested and stored at -80°C before DNA and RNA extraction for genotype-by-sequencing (GBS), whole genome bisulfite sequencing (WGBS), whole genome sequencing (WGS), RNA sequencing (RNA-Seq), and/or small RNA sequencing (sRNA-Seq). High quality DNA was extracted using CTAB protocol (Doebley and Stec 1991) with a modification to remove RNA with RNase A and purify DNA again with phenol and chloroform. Total RNA was extracted from seedlings using a standard TRIzol protocol (Invitrogen, Carlsbad, CA).

### Phenotypic data collection and analysis

W22 and backcross lines were grown at the University of Wisconsin West Madison Agricultural Research Station, Madison, WI in summer 2014, 2016 and 2017. For each BC line and W22 used for phenotype investigation, a population of 72 seeds was planted in six blocks with one plot for each line. Each plot was 3.6 m long, 0.9 m wide and was sown with 12 seeds. All of the BC_1_ plants were evaluated for plant height (length of the primary stalk from the ground to the node of top leaf) and the height of the top ear node (height from ground to base of top ear). Advanced backcross lines (mainly BC_6_ and BC_7_, and partial for BC_8_) were evaluated for twelve traits. Three plant architecture traits were investigated: plant height, the height of top ear node, and ear barrenness. Five primary ear morphology traits were investigated: cupules per rank (number of cupules in a single rank from base to the tip of the ear), kernel row number around the ear, ear diameter, ear length, and seed set rate. Three primary tassel traits were investigated: tassel branch number, pollen-shedding or not (BC_7_ and BC_8_), and days to anthesis (BC_7_ and BC_8_). One seedling trait was investigated: primary root length (BC_8_). Student’s t-test was used for plant height, the height of first ear node, ear diameter, ear length, seed setting rate, tassel branch number and root length, which have normality distribution. Mann-Whitney-Wilcoxon Test was used for cupule per rank, kernel row number, days to anthesis, and pollen-shedding, which were not normally distributed. The seeds used for measuring primary root length were treated with fungicide, rolled up in the germination paper, and placed in an incubator in the dark at 37°C for two days. All statistical analyses were performed using the R package for Statistical Computing.

### SNP genotyping methods

96-plex libraries were constructed according to genotype-by-sequencing (GBS) protocol (Elshire *et al.* 2011). Each DNA sample was digested with ApeKI restriction endonuclease, and ligated to barcode adaptors. All samples were then pooled together for PCR to increase the fragment pool. Single-end sequencing (100 bp reads) of 96-plex library per flowcell channel was performed on the Illumina HiSeq2000/2500. On average, about 2 million reads were collected from each sample, resulting in roughly 0.1X coverage of the maize genome (0.2X to 0.3X after accounting for two or three replicates). The resulting Illumina FASTQ files were processed with the TASSEL-GBS pipeline for SNP calling (Glaubitz *et al.* 2014). The reads were trimmed and aligned to the B73 reference genome (AGPv2). Polymorphisms were called under ZeaGBSv2.7 Build, which contained 955,690 SNPs derived from more than 60,000 samples (http://www.panzea.org/).

### Fluorescence in situ hybridization (FISH)

For germination and root tip harvest, captan-treated seeds were germinated at 28°C in moist vermiculite. After three or four days, the distal 1-1.5 cm segment was harvested from primary roots 3.5-5.5 cm in length. Excised root tips were transferred immediately to a humidified, 0.6-mL microcentrifuge tube with a hole in the lid and treated with nitrous oxide gas (160 psi, 2.5 hr) to stop development at metaphase (Kato 1999). Roots were subsequently fixed for 10-12 min in 90% ice-cold acetic acid, rinsed twice with cold 70% ethanol, transferred to a new tube of 70% ethanol, and stored at -20°C. Because the primary roots from Bravo and Blanco teosinte lines were thin and produced few metaphase spreads, in a subsequent planting, the germinated seeds were transferred to six-packs containing Pro-Mix BX and allowed to grow for about 13 days before processing the roots as described above. A balanced fertilizer and a water-soluble iron chelate were applied during the growth period.

The chromosome morphology of two or three roots from each seed stock was examined using FISH (Kato *et al.* 2004). The Cent4 probe used was re-designed to remove homology to 180-bp knob heterochromatin (Lamb *et al.* 2007). Probe concentrations were as described in Birchler *et al.* (2007), with the exception that the blue and green probes were labeled with coumarin-5-dUTP (custom synthesis, Perkin Elmer Life and Analytical Sciences) and fluorescein-12-dUTP (Perkin Elmer), respectively. Detailed protocols are available in (Kato *et al.* 2011). Images were acquired using an Olympus BX61 fluorescence microscope fitted with a Cool-1300QS CCD camera (VDS Vosskühler) and FISHView EXPO 4.5 software (Applied Spectral Imaging). Image background was increased or decreased using the Curves function of Adobe Photoshop CS3 while maintaining original signal strength as much as possible.

### WGS data generation, ***data processing***, ***and alignments***

DNA concentration was verified using the Qubit® dsDNA HS Assay Kit (Life Technologies, Grand Island, NY). Samples were prepared according the TruSeq Nano DNA LT Library Prep Kit (Illumina Inc., San Diego, CA, USA) with minor modifications. Samples were sheared using a Covaris M220 Ultrasonicator (Covaris Inc, Woburn, MA, USA), and were size selected for an average insert size of 350 bp using SPRI bead-based size exclusion. Quality and quantity of the finished libraries were assessed using an Agilent DNA1000 chip and Qubit® dsDNA HS Assay Kit, respectively. Seven libraries were normalized to 2 nM and then pooled into one lane. Cluster generation was performed using the Illumina Rapid PE Cluster Kits v2 and the Illumina cBot. Single-read, 100 bp sequencing was performed, using Rapid v2 SBS chemistry on an Illumina HiSeq2500 sequencer. All reads were trimmed with Cutadapt/1.8.1 (Martin 2011) to remove the Illumina TruSeq Universal adapted as well as requiring a minimum read length of 30 and a phred score of 10. Reads were aligned to the W22 maize reference genome and the *de novo* transcript assemblies using Bowtie2/2.2.4 (Langmead and Salzberg 2012). Uniquely mapping reads were further processed into 100bp windows across the maize genome for each sample and counts per million were calculated. CNVseq (Xie and Tammi 2009) was used to identify regions of copy number variation between the Bravo BC samples and the W22 sample. The BravoBC_1_ and BravoBC_6_ bam files containing uniquely aligning reads were merged to generate a single BravoBC bam file. This BravoBC file was used as the test sample in CNVseq with W22 as the reference sample. CNVseq input criteria used included a log2 cutoff of 2, p-value cutoff of 0.001, and a window size of 1kb. CNVseq output was further filtered to include only those regions with a log2FC > 3 resulting in 436 unique regions. HTseq/0.5.3 (Anders *et al.* 2015) was used in order to count the number of reads mapping to these specific regions across all samples. Additional filtering included the removal of regions containing insufficient coverage across all non-Bravo BC samples as well as a calculated log2FC value for Bravo BC vs Blanco BC_1_, BlancoBC_6_, and Blanco Teosinte > 3 resulting in 19 unique regions.

### RNA-seq data generation, ***data processing alignment***, ***and analysis***

RNA was quantified and quality tested by NanoDrop and Agilent RNA NanoChip. Samples were prepared according to the TruSeq Stranded mRNA Library Prep (Illumina, Inc., San Diego, CA, USA). One ug of RNA was transferred to a final volume 50ul with nuclease-free water, polyA selected and fragmented 6 minutes. First strand (with random hexamers) and second strand cDNA and adenylate 3’ ends were synthesized, universal and multiple indexing adapters (contain unique 6-bp indices barcode sequences) were ligated to the ends of the double-stranded cDNA, and the DNA fragments were enriched by 11 cycles of PCR-based on the adapter sequences primers. Quality and quantity of the finished libraries were assessed using an Agilent DNA1000 chip and Qubit® dsDNA HS Assay Kit, respectively. Eight indexed DNA libraries are normalized to 10 nM and then pooled into one lane of an Illumina HiSeq2000/2500 sequencer (100 bp, single end). Approximately 20 million reads were generated for each sample (Table S1). Raw reads were trimmed using cutadapt version 8.1.1 using the -m 30 -q 10 -- quality-base=33 options. All reads were aligned to W22 genome sequence (Springer *et al.* 2018) or the *de novo* transcript assemblies using Tophat2 (Kim *et al.* 2015). Four mismatches, a minimum intron size of 5 bp and a maximum intron size of 60,000 bp were used for alignment. More than 80% of reads were mapped on to the reference (Table S1). Transcript quantification was performed with HTSeq-count (Anders *et al.* 2015), using the maize W22 gene annotation. PCA analysis was performed with the plotPCA function (Wickham 2016) using the variance stabilizing transformation (VST) matrix for RNA counts which were calculated by the R/Bioconductor package DESeq2 (Love *et al.* 2014). Differentially expressed genes were identified using DESeq2 and filtered using a fold-change cutoff of 2 and an FDR-adjusted p-value cutoff of < 0.05. TE family expression was quantified as in (Anderson *et al.* 2018). Briefly, the W22 TE annotation file was modified to resolve nested TEs using RTrackLayer (Lawrence *et al.* 2009) and used as input to HTSeq-count. The SAM output was then parsed using a custom script where mapped reads are assigned to TE families when they mapped uniquely to a single TE or when multi-mapped but hit only a single TE family. Differential expression analysis was performed using DESeq2, using the same contrasts and cutoffs as used to call DE genes.

### RNA-seq de novo transcriptome assembly

A *de novo* assembly of the Bravo backcross transcriptome was generated using 52 M reads concatenated from 3 Bravo backcross RNA-seq libraries (Table S1). Illumina TruSeq adapters were removed with cutadapt v1.7.1 (Martin 2011) and only reads >= 70 bp retained. Reads were then filtered with FASTX-Toolkit (http://hannonlab.cshl.edu/fastx_toolkit/index.html) v0.0.14 *fastx_artifacts_filter* and *fastq_quality_trimmer* to remove artifacts and retain only reads Q33 or greater; final library quality was assessed with fastqc v0.11.7. *De novo* assembly was performed with Trinity (Grabherr *et al.* 2011) version r20140717 with a --min_contig_length of 200bp. Relative expression of each ‘gene’ isoform was estimated with RSEM v1.3.0 (using bowtie2 v2.3.0 for alignment) using the Trinity utility script *align_and_estimate_abundance.pl* from Trinity version r20140717. For differential expression testing a SuperTranscript reference (Davidson *et al.* 2017) was generated using the Trinity utility script (Haas *et al.* 2013b) *Trinity_gene_splice_modeler.py* from Trinity version v2.6.6. Differentially expressed transcripts were identified by mapping RNAseq reads to the transcriptome assembly using hisat2 (Kim *et al.* 2015) v2.1.0, retaining unique-mapping reads, and running DEseq2. The differentially expressed *de novo* transcripts were then analyzed for putative coding regions using TransDecoder-v5.3.0 by retaining ORFs >= 100 amino acids or regions with homology to protein domains. Protein homology was identified using blast+ (Camacho *et al.* 2009) v2.7.1 *blastp* against ether Swissprot (ftp://ftp.ncbi.nlm.nih.gov/blast/db/swissprot.tar.gz downloaded 31 May 2018) or UniRef90 (ftp://ftp.uniprot.org/pub/databases/uniprot/uniref/uniref90/uniref90.fasta.gz downloaded 31 May 2018) databases and using HMMER v 3.1b2 (Johnson *et al.* 2010) *hmmscan* to search the Pfam-A database (ftp://ftp.ebi.ac.uk/pub/databases/Pfam/current_release/Pfam-A.hmm.gz downloaded 31 May 2018).

### Small RNA-seq data generation, ***data processing alignment***, ***and analysis***

Samples were prepared according to the TruSeq Small RNA Library Prep Kit (Illumina, Inc., San Diego, CA, USA). One ug of RNA was transferred to a final volume 6ul with nuclease-free water. The 3’ (containing the unique six-base indexes barcode sequences) and 5’ adapters were ligated, and the reverse transcription was performed followed by amplification (11 cycles). PCR cDNA products were further purified using 6% TBE Gel and gel bands corresponding to molecular sizes of 145-160bp were excised. A total of 24 small RNA libraries (Table S1) were pooled into one lane of Illumina HiSeq2500 sequencer for sequencing (single-end reads of 50-bp) at University of Wisconsin (Madison, WI). Raw reads were first de-multiplexed and passed to FastQC for initial quality control. Reads from small-RNA sequencing contain the 3’ sequencing adapter because the read is longer than the molecule that is sequenced. Thus, the high-quality clean reads were processed using cutadapt v1.15 (Martin 2011) to remove the 3’ adapter sequence (TGGAATTCTCGGGTGCCAAGG). Any reads that did not contain an adapter were discarded, and only 18∼42-nt long reads were retained for subsequent analyses. Adaptor-free reads from rRNA, tRNA, snoRNA, and snRNA fragments were removed by aligning them against the corresponding genomic sequences of *Zea mays* genome using Bowtie v1.1.2 (Langmead *et al.* 2009) allowing for two mismatches. The remaining reads were mapped to the W22 (https://www.maizegdb.org/genome/genome_assembly/Zm-W22-REFERENCE-NRGENE-2.0) and B73 genomes (ftp://ftp.gramene.org/pub/gramene/release-58/fasta/zea_mays/dna/) for sRNA samples from Doebley and Flint-Garcia Laboratories, respectively. At this step, ShortStack v3.4 was employed with parameters ‘-bowtie_m 1000 -ranmax 50 -mmap u -mismatches 0’, which employed local genomic context to guide decisions on proper placements of multi-mapped sRNA-seq reads (Johnson *et al.* 2016). To calculate and compare small RNA abundance in different samples, the genome was tiled into 500-bp windows and reads whose 5’ end nucleotides fall within a window were counted. sRNA with a size between 19 and 26 nucleotides were selected and sRNA abundance for each window were calculated as reads per million (RPM) of total reads. Differential comparisons of expression abundance were conducted by the R package ‘DESeq2’ (Love *et al.* 2014).

### WGBS data generation, data processing alignment and analysis

Genomic DNA was sheared to 200-300bp fragments, which were then subject to end repair, A-tailing, adapter ligation and dual-SPRI size selection using KAPA library preparation kit (KK8232) following manufacturer’s instructions. The size (250bp – 450bp) selected library was then treated with bisulfite to convert unmethylated cytosine to uracil using Zymo EZ DNA methylation lightning kit (D5031). The converted DNA was then PCR amplified using KAPA HiFi HotStart Uracil + (KK2801) with the following program: 95°C/2min, 8 times of 98°C/30s, 60°C/30s, 72°C/4min, and a final extension at 72°C for 10min. To increase PCR efficiency, the bisulfite converted DNA was split into two equal parts and two parallel PCR reactions were performed, the final PCR products were combined and cleaned up together using SPRI beads. Library quality was checked using Agilent Bioanalyzer to ensure that the libraries size is in the right range (200-700bp with a peak around 300bp). Library quantity was checked using Picogreen to make sure that the final concentration is >2nM. Libraries that passed quality control were then equally pooled and sequenced across multiple lanes on an Illumina HiSeq2500 machine. For each library, 150M to 252M paired-end 125bp reads were generated (Table S1). Adapter sequences were trimmed and read quality was assessed using Trim Galore! with the default parameters and paired-end reads mode. Reads that passed quality control were mapped to maize W22 genome using BSMAP (Xi and Li 2009), allowing at most 5 mismatches. Nucleotides with a quality score less than 20 were trimmed from 3’ end of reads. Reads that are uniquely mapped and that are properly paired were used to extract methylation status at individual cytosine using methratio.py which is included in the BSMAP package. At this stage, duplicate reads due to PCR bias were removed. Also, nucleotides in the overlapped part of paired hits are only counted once instead of twice. The final output of methratio.py contains the number of methylated and unmethylated reads for each cytosine. This file was used to create 100bp non-overlapping sliding windows across the maize chromosomes for each of the three cytosine contexts, CG, CHG and CHH (H = A, C or T). Within each 100bp window and for each sequence context, the total number of methylated/unmethylated reads for each cytosine was summed and used to derive methylation levels of the 100bp window, (#C/(#C+#T)). Differential methylation region (DMR) calling was performed between the Bravo BC lines (BravoBC_1_ and BravoBC_6_) and W22 to identify context-specific DMRs. We required CG/CHG DMRs to have a minimum of 3 symmetrical CG/CHG sites (6 Cs total for the two strands) and a minimum methylation difference of 60% between the two samples. DMRs were called based on 100bp bins with >2x coverage in both samples. Coverage was defined as the ratio of the total number of times a cytosine was covered by the total number of cytosines of that specific context.

For Principal Components analysis, the average DNA methylation level was determined for 100bp non-overlapping windows for all three sequence contexts (CG, CHG, and CHH). The matrix of CG DNA methylation levels for each 100bp window for the seven genotypes was used to perform a principle components analysis (R package prcomp).

## Data availability

The WGBS data are available under the National Center for Biotechnology Information Bioproject accession PRJNA526266; the RNA-Seq data are available under PRJNA528342; the WGS data is available under PRJNA528290, and the sRNA-seq data is available under PRJNA528352. The Sequence Read Archive (SRA) accession numbers are available in Table S1. The GBS data reported in this paper have been deposited in the figshare database (https://figshare.com/s/a882aff235818fc1762c).

## Acknowledgements

We thank Xuehua Zhong and Chin Jian Yang for helpful discussions regarding this work and Jonathan Giesler for technical assistance. We thank the University of Wisconsin Biotechnology Center DNA Sequencing Facility for providing next generation sequencing consultation and services. This research was supported by the National Science Foundation (NSF) grants IOS 1238014 (J.F.D. and S. F.-G.), IOS-1237931 (to N.M.S) and IOS-1444514 (J.A.B.), and by USDA-NIFA grant 2016-67013-24747 (to N.M.S), and by USDA-ARS base funds (to S. F-G).

